# Gestational Cd Exposure in the CD-1 Mouse Sex-Specifically Disrupts Essential Metal Ion Homeostasis

**DOI:** 10.1101/2021.11.06.467551

**Authors:** Thomas W. Jackson, Oliver Baars, Scott M. Belcher

**Author notes:** Corresponding Author: Scott M. Belcher. Email Address: Thomas W. Jackson Oliver Baars Scott M. Belcher.

## Abstract

In CD-1 mice, gestational-only exposure to cadmium (Cd) causes female-specific hepatic insulin resistance, metabolic disruption, and obesity. To evaluate whether sex differences in cadmium uptake and changes in essential metal concentrations contribute to metabolic outcomes, placental and liver cadmium and essential metal concentrations were quantified in male and female offspring perinatally exposed to 500 ppb CdCl_2_. Exposure resulted in increased maternal liver Cd^+2^ concentrations (364 μg/kg) similar to concentrations found in non-occupationally exposed human liver. At gestational day (GD) 18, placental cadmium and manganese concentrations were significantly increased in exposed males and females, and zinc was significantly decreased in females. Placental efficiency was significantly decreased in GD18 exposed males. Increases in hepatic Cd concentrations and a transient prenatal increase in zinc were observed in exposed female liver. Fetal and adult liver iron concentrations were decreased in both sexes, and decreases in hepatic zinc, iron, and manganese were observed in exposed females. Analysis of GD18 placental and liver metallothionein mRNA expression revealed significant Cd-induced upregulation of placental metallothionein in both sexes, and a significant decrease in fetal hepatic metallothionein in exposed females. In placenta, expression of metal ion transporters responsible for metal ion uptake was increased in exposed females. In liver of exposed adult female offspring, expression of the divalent cation importer (*Slc39a14/Zip14)* decreased, whereas expression of the primary exporter (*Slc30a10*) increased. These findings demonstrate that Cd can preferentially cross the female placenta, accumulate in the liver, and cause lifelong dysregulation of metal ion concentrations associated with metabolic disruption.

## 1. Introduction

Cadmium (Cd) is a ubiquitous environmental contaminant ranked 7^th^ on the list of toxicants of concern by the Agency for Toxic Substances and Disease Registry (ATSDR 2012 Sep). In human adults, elevated Cd exposure increases risk of cardiovascular disease, hypertension, diabetes, osteoporosis, impaired kidney function, and cancer (Tellez-Plaza et al. 2013; Åkesson et al. 2014). Emerging data suggests developmental Cd exposure increases risk for childhood obesity (King et al. 2015; Lamas et al. 2016). Using a mouse model of human gestational Cd exposure, we previously demonstrated that Cd body burdens equivalent to women of childbearing age sex-specifically caused hepatic steatosis, dyslipidemia, glucose intolerance, insulin resistance, and obesity in female offspring (Jackson et al. 2020). The resulting later-in-life obesogenic effects of gestational Cd exposure in females were found to be mediated by well-established mechanisms involved in cumulative Cd toxicity (Sabolić et al. 2010; Nair et al. 2013; Nemmiche 2016 Nov 1). Specifically, oxidative stress, endoplasmic reticulum stress, and mitochondrial dysfunction were notable at birth in exposed female offspring and progressed to pathologic metabolic disease without continued Cd exposure. The resulting severe liver damage and obesity were linked with female-specific developmental disruption of hepatic retinoic acid signaling, programing of hepatic toxicity, disrupted insulin signaling, metabolic dysregulation leading to obesity, and an increase biomarkers of hepatocellular carcinoma (Jackson et al. 2020).

In apparent contrast to our findings that brief developmental Cd exposures caused adverse metabolic impacts later in life, Cd is generally considered a cumulative toxicant, meaning that toxic effects develop over time as Cd accumulates due to slow elimination (Waalkes 2003; Jacobo-Estrada et al. 2017). Previous studies examining circulating maternal Cd transfer to the fetus found that Cd is largely sequestered in placenta, resulting in little or no fetal Cd accumulation (Mikolić et al. 2015). However, those conclusions are from studies that compared placental Cd concentrations to Cd in whole fetus, an approach that underestimates Cd sequestered in target tissues such as fetal liver (Sigel et al. 2013; Mikolić et al. 2015). Further, impacts of relatively low concentrations of Cd, differences in Cd exposure in male and female fetus, and impacts of fetal sex as a modifier of placental ability to sequester Cd are not well studied.

Cadmium accumulates primarily in liver and kidney, with much lower levels accumulating in pancreas, heart, testis, bone, and neural tissues (ATSDR 2012 Sep). Organ-specific patterns of Cd uptake and toxicity are mediated by differential expression of SLC39 transporters, SLC30 transporters, and divalent metal transporter-1 (DMT1) (Fujishiro et al. 2012). The functional roles of Zip8 (SLC39A8) and Zip14 (SLC39A14) Zinc (Zn) and Manganese (Mn) transporter proteins in regulating absorption, intracellular uptake, accumulation, and toxicity of Cd in testis, kidney, and liver are well established (Sabolić et al. 2010; Fujishiro et al. 2012; Aydemir and Cousins 2018; Fujishiro and Himeno 2019). The physiological role of divalent metal ion transporters is to regulate essential divalent metal ion homeostasis, a regulatory process adversely impacted by Cd (Aydemir and Cousins 2018). In addition to Zn and Cd ion transport, Zip14 also functions as the liver’s Mn transporter, which works with Mn efflux transporter Slc30a10 (ZnT10) to regulate Mn homeostasis in various tissues including liver (Mercadante et al. 2019: 10).

The placenta is a critical regulator of offspring health that coordinates fetal nutrition and oxygen transfer, and responds to Cd exposure by sequestering Cd and upregulating expression of metallothionein (Lau et al. 1998; Everson et al. 2019). During pregnancy, Zip14, Slc39a4 (Zip4), DMT1, and Slc30a2 (ZnT2) facilitate uptake and transfer of essential metals and Cd from maternal blood vessels to placenta, and subsequent transfer from placenta to fetal cord blood (Espart et al. 2018). There is ample evidence demonstrating that Cd alters homeostasis of Zn, Fe, and Mn (Moulis 2010). For example, adult Cd exposure induces anemia by altering iron metabolism and homeostasis, whereas increased pregnancy and lactation Cd levels associated with Fe deficiency (Akesson et al. 2002; Horiguchi et al. 2011). We also observed anemia at birth in newborn mice following maternal exposure to 500 ppb CdCl_2_, a find that suggests iron concentrations were altered by gestational exposure to relatively low Cd concentrations (Jackson et al. 2020).

Considering the important roles of metallothioneins and metal ion transporters in Cd toxicity and essential metal homeostasis, we hypothesized that gestational Cd exposure alters placental metallothionein and dysregulates Zn transporter expression in female placenta and Cd accumulation in female fetal livers. Further, increases of Cd in female offspring liver would result in lifelong changes in metallothionein and hepatic transporter expression, and dysregulated adult hepatic divalent cation concentrations. To evaluate these hypotheses, inductively coupled plasma mass spectrometry was used to quantify concentrations of Cd^2+^, Fe^2+^, Mn^2+^and Zn^2+^in maternal livers, placenta, and offspring livers following perinatal exposure of pregnant dams to 500 ppb CdCl_2_ in drinking water. Impacts of exposure on transcription of genes involved in metal ion homeostasis, and epigenetic regulation of those genes, were evaluated. Analyses included samples collected from gestational day (GD)18 through postnatal day (PND)120 to describe progression of metal homeostasis disruption and evaluate differences in hepatic Cd concentrations in male and female offspring following gestational Cd exposure.

## 2. Materials and Methods

### 2.1 Animal Husbandry

All animal procedures were carried out as previously described (Jackson et al. 2020) following recommendations of the Panel on Euthanasia of the American Veterinary Medical Association, and were approved by the North Carolina State University Institutional Animal Care and Use Committee. Study animals were housed in single-use polyethylene cages (Innovive, San Diego, CA) with Sanichip bedding (PJ Murphy Forest Products Corp, Montville, NJ) and pulped virgin cotton fiber nestlets (Ancare, Bellmore, NY) on a 12:12 light cycle at 25°C and 45%-60% average relative humidity in an AAALAC accredited animal facility. Defined AIN-93G diet (D10012G; lot: 17010510A6, Research Diets, New Brunswick, NJ) and sterile drinking water produced from a reverse osmosis water purification system (Millipore Rios with ELIX UV/Progard 2, Billerica, MA) was supplied *ad libitum*. Strain CRL:CD-1(ICR) ;CD-1) male and female breeder mice were obtained from Charles River Laboratories (Raleigh, NC) and assigned a randomized identification number. Beginning 2 weeks prior to mating and ending on postnatal day 10 (PND10), drinking water of dams in the Cd-exposed group was supplemented with a final concentration of 0.5 μg/L (500 ppb) cadmium chloride (CdCl_2_; CAS 10108-64-2; 99.99% purity, Lot MKBM1769V, Sigma Aldrich). Beginning on PND10, drinking water for both control and exposed study groups for the remainder of the study was Cd-free.

A subset of dams (n=3 per group) was euthanized by CO_2_ asphyxiation and rapid decapitation at GD18 and fetuses isolated by dissection. Those procedures were completed 2-3 hours after lights on. Dams and fetuses were weighed, and tissues were collected. The intrauterine position of each fetus was recorded and fetal genetic sex was determined by Sry-specific PCR as previously described (Lambert et al. 2000). For the remaining litters, litter size, pup sex, and pup weight were recorded on the day following parturition (PND1). Offspring were separated from dams at PND21 and housed 2-4 per cage separated by sex and study group. From each litter, one male and one female representative of the mean litter body weight was euthanized at PND1, PND21, PND42, PND90, and at study termination on PND120. Study animals remained in their home cage, with food and water available, until euthanized by carbon dioxide asphyxiation or transcardiac perfusion under isoflurane anesthesia.

All study animals were necropsied with study samples and tissues isolated at the time of sacrifice. Systematic bias was avoided by housing animals randomly on cage racks and ensuring the timing and order of measurements, data collection, and experimental manipulations were the same. Necropsy was performed at the same approximate time of day, independent of study group, and dependent on date of birth. For adult females, analysis and sample collection were performed in estrus. All manipulations, tissue collections, and analysis were done by investigators blinded to study group.

### 2.2 Tissue Isolation and ICP-MS

At necropsy, tissues were dissected, frozen on powdered dry ice, and stored at -80°C until prepared for analysis. Samples were digested in 1 mL trace metal grade nitric acid (Thermo Fisher, Waltham, MA; Catalog #A509) and 1 mL hydrogen peroxide (Thermo Fisher, Waltham, MA; Catalog #H341) at 100 °C for 3 hours. Digested samples were allowed to cool and filtered through a 0.22-μm filter (Millipore Sigma, Burlington, MA; Catalog #SLMP025SS). The filtered solution was transferred to a 50 mL conical tube, internal standards were added, samples were then adjusted to a final volume of 50 mL with deionized water and vortexed for 30 seconds. Identically prepared samples lacking added tissue were used as procedural blanks. Tissue metal concentrations determined using an ICAP RQ ICP-MS (Thermo Fisher, Waltham, MA) at the NCSU Molecular Education, Technology and Research Innovation Center. Bovine liver standard (SRM 1577c) was purchased from the National Institute of Science and Technology. The accuracy and precision of our analysis for tissue cadmium, zinc, iron, and manganese levels were assessed with a standard reference bovine liver () in each analysis. Certified values for the metals in the reference bovine liver were 97 μg/kg for Cd and 181, 198, and 10.5 μg/g dry-tissue weight for Zn, Fe, and Mn, respectively. Observed mean values SRM 1577c (n = 17) of 98 μg/kg for Cd and 189, and 206, and 13 μg/g dry-tissue weight for Zn, Fe, and Mn, respectively. The coefficient of variation was 7.0% for Cd, 12.1% for Zn, 13.3% for Fe, and 13.8% for Mn. Limits of detection (LOD) were 0.1 ppb for each analyte. Values below the detection limit were replaced by interpolated values calculated by dividing the LOD by the square root of 2 (Sanford et al. 1993).

### 2.3 Quantitative RT-PCR analysis

Total RNA was isolated using the RNEasy Mini Kit (Qiagen, Valencia, CA). One μg of RNA was reverse transcribed using the high-capacity cDNA reverse transcription kit (Applied Biosystems; Grand Island, NY) following manufacturer’s protocols. Standard PCR amplification was performed in triplicate on a Step One Plus Real-Time PCR System (Applied Biosystems; Grand Island, NY) in a final volume of 20 μL containing ∼10 ng of cDNA (1.5 μL of RT product), 1x Universal Master Mix and TaqMan expression assay primers (Supplemental Table 1); Applied Biosystems; Grand Island, NY). Relative expression was quantified using the 2^ΔΔCt^ method, in which ΔΔCt is the normalized value.

### 2.4 Analysis of DNA methylation in putative promoter regions

Bisulfite pyrosequencing assays were developed to quantitatively measure the level of methylation at CpG sites within putative promoter regions upstream of the metal ion transporters Slc30a10 and Slc39a14. Genomic DNA (1 μg) was treated with sodium bisulfite using the EZ DNA Methylation Gold Kit per manufacturer’s instructions (Zymo Research; Irvina, CA). Bisulfite converted DNA (∼20 ng) was amplified by PCR in a 25 μl reaction volume using HotStartTaq plus DNA polymerase (Qiagen; Germantown, MD) with 1.5mM MgCl2 and 0.12 μM each of the forward and reverse primer (Supplemental Table 2), and 2.5 μl of CoralLoad Concentrate (Qiagen). The reverse primer of each pair was biotin conjugated at the 5’ end, with single stranded amplicons isolated on the Pyrosequencing Work Station and pyrosequencing was performed on a Pyromark Q96 MD instrument (Qiagen; Germantown, MD). Pyrosequencing assays were performed in duplicate, and values reported as the mean methylation of CpG sites contained within the sequence analyzed. Using those methods, a minimum 5% difference in methylation can be detected (Murphy et al. 2012).

### 2.5 Data and Statistical Analysis

All procedures, measurements and endpoint assessments were made by investagators blinded to treatments, litter, and sex when appropriate. To avoid influence of extreme litter size on endpoint sensitivity, litters with fewer than 6 pups were excluded from analysis and litters with greater than 14 pups were culled to a maximum of 12 (Palmer and Ulbrich 1997). Analysis of body weight, placenta weight, and placental efficiency, data were analyzed using two-way ANOVA (sex, exposure), with litter size included as a covariate. Trace metal concentration data were analyzed using a two-way ANOVA (sex, exposure) for each metal at each timepoint with litter size and litter included as covariates. If overall effects were significant, a Tukey’s least significant differences post hoc test was performed to evaluate pair-wise differences. Effect sizes were calculated depending on the statistical test, values for “η^2^” for ANOVA, “Cohen’s d” for t-tests, and “r” for Mann-Whitney U are reported. Significance between differences in values were defined as *p* < .05. All data was analyzed using Prism^®^v9 (GraphPad; La Jolla, California) and SPSS V.26 (IBM, California). A table of all statistical analyses and results for morphometrics and metal concentrations is included in Supplemental Table 3.

## 3. Results

### 3.1 Effects of gestational Cd exposure on body weight and placental efficiency

Placenta weight of Cd-exposed males was increased 18% from 0.11 ± 0.17 to 0.13 ± 0.39 grams (p = .02; Figure 1A) and placental efficiency was decreased 14% from 14 ± 2 to 12 ± 4 (p = .04; Figure 1B). Placenta weight and placental efficiency were unchanged in Cd-exposed females.

**Figure 1:**
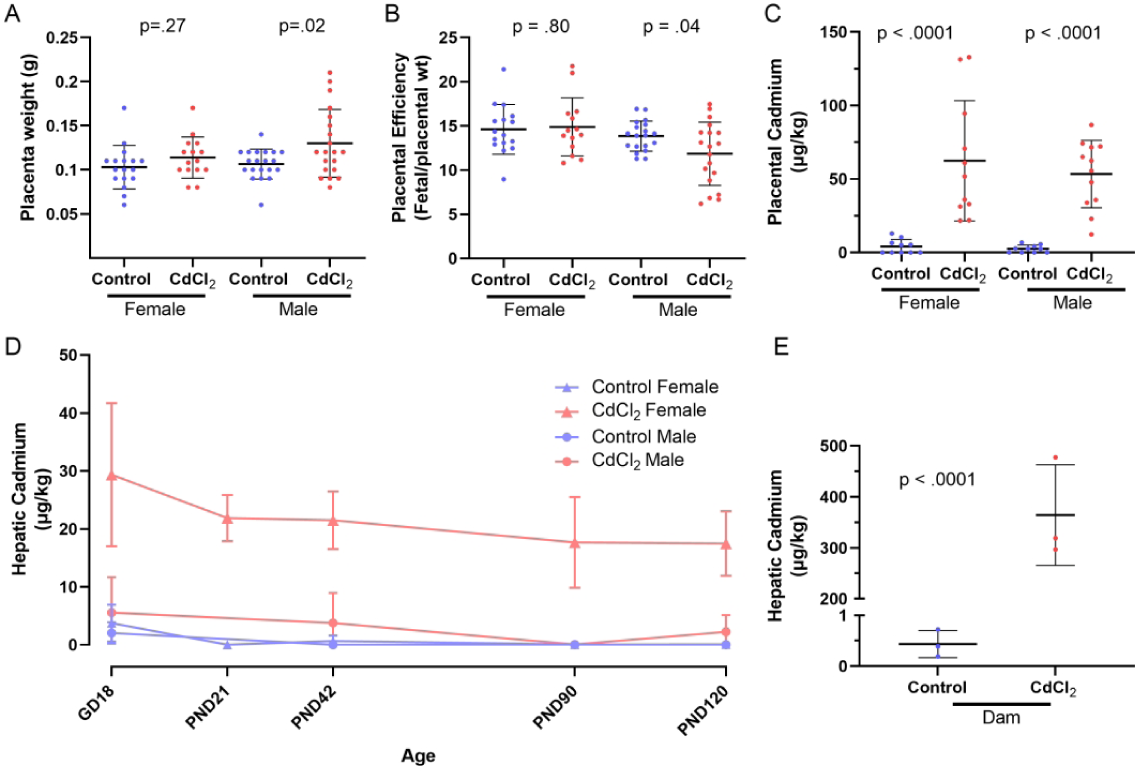
Effects of gestational CdCl2 exposure on placental efficiency and tissue cadmium levels. At GD18, placentas of female fetuses were unaffected by exposure to gestational CdCl2, whereas male placenta weights were increased (A). Placental efficiency is shown as fetal weight divided by placental weight, where female offspring show no effect of gestational CdCl2 exposure and male placental efficiency is reduced (B). Cadmium concentration of tissues were quantified using ICP-MS. In the placenta of offspring exposed to CdCl2 during gestation, both sexes showed accumulation with an average in females of 62 μg/kg and in males of 53 μg/kg (C). Offspring liver concentrations of Cd are shown plotted against age (D). In offspring female livers at GD18 following gestational exposure to CdCl2, a significant accumulation of Cadmium was noted with an average of 29 μg/kg, whereas males did not have a significantly elevated accumulation of Cd with an average of 5.6 μg/kg. An average of 22 μg/kg Cd was detected in females gestationally exposed to CdCl2 in livers at PND21, 21 μg/kg Cd in livers at PND42, 18 μg/kg Cd in livers at PND90 and 17 μg/kg Cd in livers at PND120. An average of 3.8 μg/kg Cd was detected in males gestationally exposed to CdCl2 in livers at PND42, 0.05 μg/kg Cd in livers at PND90, and 2.3 μg/kg Cd in livers at PND120. In livers isolated from dams exposed to CdCl2 at GD18 of their pregnancy, an average of 364 μg/kg Cadmium was detected (E). Placenta weight and efficiency: Females: control, n=17; CdCl2, n=14. Males: control, n=20; CdCl2, n=22. Cadmium levels: GD18: Females: control, n=9; CdCl2, n=11. Males: control, n=8; CdCl2, n=10. PND21: Females: control, n=4; CdCl2, n=4. PND42: Females: control, n=5; CdCl2, n=4. Males: control, n=4, CdCl2, n=4. PND90: Females: control, n=6; CdCl2, n=4. Males: control, n=4; CdCl2, n=6. PND120: Females: control, n=5, CdCl2, n=4. Males: Control, n=3; CdCl2, n=5. Dams: control, n=3; CdCl2, n=3. Samples were collected from 3 litters per treatment. All values shown are mean ± SD. The level of statistical significance for differences between mean values of control and CdCl2-exposed groups was determined by a two-way ANOVA (treatment, sex) with a Tukey’s post hoc test for all experiments and is indicated by * (p<0.05). Litter was included as a covariate.

### 3.2 Prenatal metal content in liver and placenta

At GD18, maternal Cd-exposure resulted in increased concentrations in mean female and male placental Cd (p < .0001; Figure 1C; Table 1). Offspring liver Cd concentrations were significantly increased in exposed females (p < .0001) but not male (p = .34; Figure 1D; Table 1). Pregnant female liver Cd concentrations were also significantly increased by exposure (p = .003; Figure 1E; Table 1). Concentrations of Zn in the placenta of female offspring were significantly decreased (p < .0001) whereas Mn concentrations were increased (p = .03). In exposed male placenta mean Mn concentrations were increased by exposure (p = .005), however there was no change in Zn concentrations (p = .78). Placental iron concentrations were not changed by Cd exposure in either sex (Table 1).

**Table 1.** Metal content of dam liver and offspring liver and placenta at GD18 measured by ICP-MS.

|  | Dam |  | Placenta |  |  |  | Liver |  |  |  |
| --- | --- | --- | --- | --- | --- | --- | --- | --- | --- | --- |
|  |  |  | Female |  | Male |  | Female |  | Male |  |
|  | Control | CdCl<sub>2</sub> | Control | CdCl<sub>2</sub> | Control | CdCl<sub>2</sub> | Control | CdCl<sub>2</sub> | Control | CdCl<sub>2</sub> |
| Cadmium (µg/kg) | 0.43 ± 0.27 | **364 ± 99\*** | 4.1 ± 4.9 | **62 ± 41** | 2.5 ± 2.5 | **53 ± 23** | 3.7 ± 3.2 | **29 ± 12** | 2.0 ± 1.9 | 5.5 ± 6.1 |
| Zinc (µg/g) | 155 ± 4.5 | 150 ± 5.9 | 157 ± 11 | **101 ± 10** | 160 ± 11 | 158 ± 8.6 | 211 ± 17 | **266 ± 25** | 190 ± 11 | 185 ± 22 |
| Iron (µg/g) | 307 ± 28 | 302 ± 8.7 | 132 ± 13 | 127 ± 13 | 137 ± 26 | 135 ± 12 | 291 ± 24 | **271 ± 24** | 311 ± 52 | **264 ± 41** |
| Manganese (µg/g) | 4.4 ± 0.13 | 4.4 ± 0.78 | 2.4 ± 0.43 | **3.1 ± 0.87** | 2.5 ± 0.46 | **3.3 ± 0.72** | 6.3 ± 1.6 | 6.6 ± 1.7 | 5.4 ± 0.86 | 5.6 ± 1.1 |

Values represent group mean ± SD; Bold\* indicates significant difference; $p < .05$ . CdCl<sub>2</sub> = 500ppb gestational Cadmium Chloride. Below LOD were defined as LOD/sqrt(2). Dam: n=3/group. Placenta: females: control n=10, CdCl<sub>2</sub> n=11; males: control n=10, CdCl<sub>2</sub> n=12. Fetal liver: females: control n=9, CdCl<sub>2</sub> n=11; males: control n=8, CdCl<sub>2</sub> n=10.

Essential metal concentrations in male livers at GD18 were unchanged in the maternal Cd exposure group (Table 1). Concentrations of Zn in livers of exposed females at GD18 were significantly increased (p < .0001) whereas hepatic Fe concentrations were decreased (p = .01) Manganese concentrations of female fetal livers were not changed by maternal Cd exposure. Hepatic Cd concentrations in exposed female offspring were positively associated with hepatic Zn (p = .003) and placental Cd (p = .046) concentrations, but negatively correlated with placental Zn (p = .001). Hepatic Zn levels were positively correlated with placental Cd (p = .001) and negatively correlated with placental Zn in exposed female offspring (p < .0001; Figure 2A). Liver Fe in liver of exposed male offspring was negatively correlated with placental Mn (p = .05), and placental Cd was positively correlated with placental Mn (p = .001; Figure 2B).

**Figure 2:**
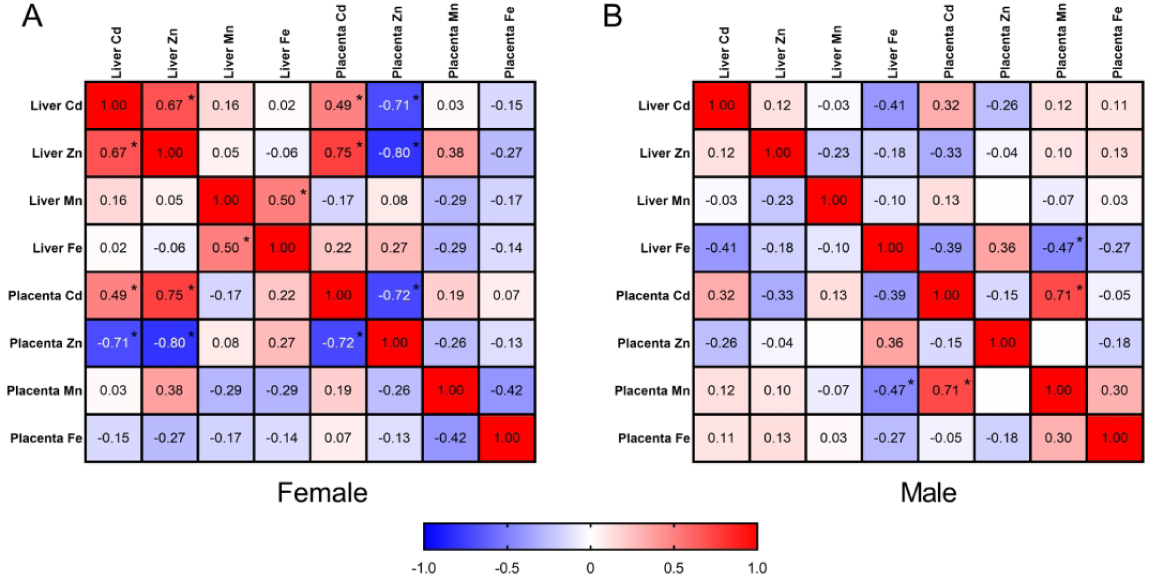
Correlation matrix of metal concentration in offspring CD-1 mouse liver and placenta at GD18. A Pearson Correlation Matrix of liver and placental metal concentration is shown for females (A) and males (B). In females, hepatic cadmium has a significant positive correlation with hepatic zinc and placental Cd, and a significant negative correlation with placental zinc. Hepatic zinc was positively correlated with placental Cd and negatively correlated with placental Zn. In males, hepatic iron was negatively correlated with placental manganese, and placental Cd was positively correlated with placental manganese. Liver: Females: control, n=9; CdCl2, n=11. Males: control, n=8; CdCl2, n=10. Placenta: Females: control, n=10; CdCl2, n=11. Males: control, n=10; CdCl2, n=12. All samples come from three unique litters per treatment.

#### 3.3 Postnatal Liver Metal Concentrations

In the postnatal liver of exposed female offspring, Cd concentrations at PND21 were significantly increased and remained elevated at each adult time point analyzed (Table 2). Liver cadmium concentrations in unexposed adult male offspring were below the limit of detection (LOD = 0.1 μg/kg), with only a modest increase in liver Cd detectable in exposed males (Table 2).

**Table 2.** Metal content of offspring liver measured by ICP-MS.

|  | PND21 |  | PND42 |  |  |  | PND90 |  |  |  | PND120 |  |  |  |
| --- | --- | --- | --- | --- | --- | --- | --- | --- | --- | --- | --- | --- | --- | --- |
|  | Female |  | Female |  | Male |  | Female |  | Male |  | Female |  | Male |  |
|  | Control | CdCl<sub>2</sub> | Control | CdCl<sub>2</sub> | Control | CdCl<sub>2</sub> | Control | CdCl<sub>2</sub> | Control | CdCl<sub>2</sub> | Control | CdCl<sub>2</sub> | Control | CdCl<sub>2</sub> |
| Cadmium (µg/kg) | 0.07 ± 0 | **22 ± 4** | .68 ± .91 | **21 ± 5.0** | 0.07 ± 0 | **3.8 ± 5.2** | 0.10 ± .08 | **18 ± 7.8** | 0.07 ± 0 | 0.10 ± 0.05 | 0.12 ± .13 | **17 ± 5.6** | 0.07 ± 0 | 2.3 ± 2.9 |
| Zinc (µg/g) | 153 ± 6.9 | **119 ± 8.7** | 141 ± 11 | **116 ± 6.1** | 133 ± 12 | 130 ± 18 | 166 ± 14 | **90 ± 8.7** | 159 ± 34 | 133 ± 19 | 149 ± 13 | **91 ± 8.8** | 143 ± 19 | 126 ± 19 |
| Iron (µg/g) | 366 ± 11 | 331 ± 29 | 284 ± 9.1 | **186 ± 10** | 294 ± 13 | 279 ± 23 | 326 ± 31 | **230 ± 41** | 324 ± 34 | **256 ± 31** | 299 ± 42 | **190 ± 24** | 352 ± 61 | **288 ± 27** |
| Manganese (µg/g) | 4.7 ± 0.49 | **3.9 ± 0.21** | 5.2 ± 0.60 | **3.3 ± 0.22** | 4.4 ± 0.27 | 4.5 ± 0.84 | 4.8 ± 0.55 | **3.7 ± 0.55** | 5.3 ± 0.48 | 4.8 ± 0.62 | 4.6 ± 0.32 | **3.5 ± 0.23** | 5.1 ± 0.12 | 4.5 ± 0.98 |

Values represent group mean ± SD; Bold\* indicates significant difference; p < .05. CdCl2 = 500ppb gestational Cadmium Chloride. Below LOD were defined as LOD/sqrt(2). PND21 Liver: n=4/group. PND42 Liver: females: control n=5, CdCl2 n=4; males: control n=4, CdCl2 n=4. PND90 Liver: females: control n=6, CdCl2 n=4; males: control n=6, CdCl2 n=4. PND120 Liver: females: control n=5, CdCl2 n=4; males: control n=3, CdCl2 n=5.

Zinc concentrations in exposed female liver significantly decreased beginning at PND21 (p = .0009) and remained decreased at each adult time point (Table 2; Figure 3A). By contrast, Zn concentrations in exposed male liver were not significantly changed at each time point analyzed (Table 2; Figure 3B). Compared to control females, hepatic Fe of exposed females was decreased throughout life, and during adulthood Fe levels were significantly decreased at each adult time point (Table 2; Figure 3C). In exposed male liver, significant decreases in liver iron concentrations were also observed, with significant decreases observed at PND90 and PND120 (Table 2; Figure 3D). Manganese concentrations were decrease in exposed female livers throughout postnatal life, whereas liver Mn in males was unchanged (Table 2; Figure 3E,3F).

**Figure 3.**
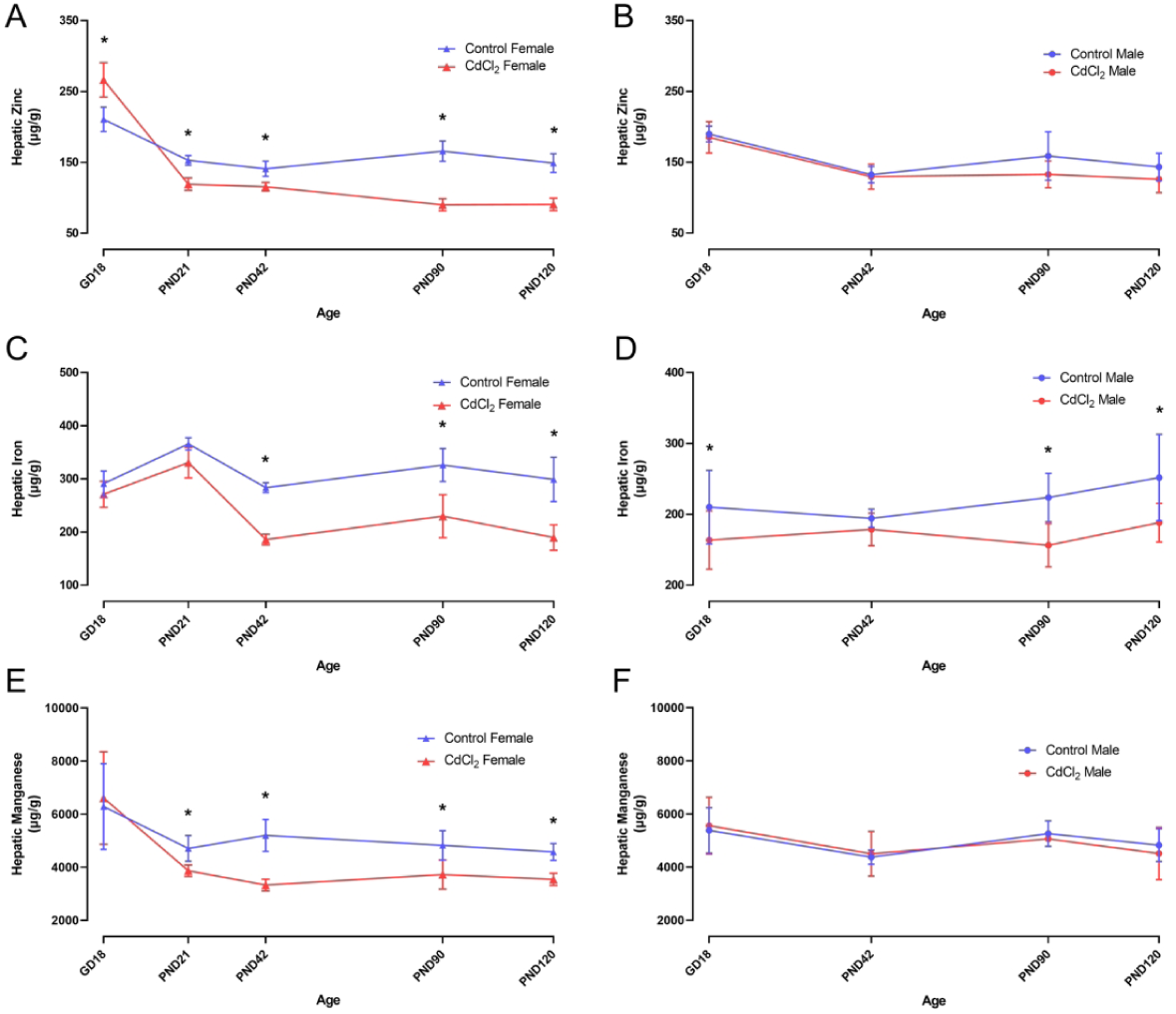
Effects of gestational exposure to CdCl_2_ on essential metal concentration in offspring CD-1 mice liver. Liver concentration of the essential metals Zinc (A, B), Iron (C, D), and Manganese (E, F) are shown for offspring female (A, C, E) and male (B, D, F) livers isolated and flash frozen at GD18, PND21, PND42, PND90, and PND120. In females, hepatic zinc at GD18 was increased from 211 in controls to 266 μg/g in Cd-exposed offspring, at PND21 was decreased from 153 to 119 μg/g in Cd-exposed offspring, at PND42 was decreased from 141 to 116 μg/g in Cd-exposed offspring, at PND90 was decreased from 166 to 90 μg/g in Cd-exposed offspring, and at PND120 was decreased from 149 to 91 μg/g in Cd-exposed offspring. In females, hepatic iron at GD18 was decreased from 291 to 271 μg/g in Cd-exposed offspring, at PND21 was unchanged, at PND42 was decreased from 284 to 186 μg/g in Cd-exposed offspring, at PND90 was decreased from 326 to 230 μg/g in Cd-exposed offspring, and at PND120 was decreased from 299 to 190 μg/g in Cd-exposed offspring. In females, hepatic manganese was unchanged at GD18, decreased from 4.7 to 3.9 μg/g in Cd-exposed offspring at PND21, decreased from 5.2 to 3.3 μg/g in Cd-exposed offspring at PND42, decreased from 4.8 to 3.7 μg/g in Cd-exposed offspring at PND90, and decreased from 4.6 to 3.5 μg/g in Cd-exposed offspring at PND120. In males, no changes were detected in essential metals at GD18 or PND42. In males at PND90 and PND120, the only changes noted in males were decreases in hepatic iron from 324 to 256 μg/g in Cd-exposed offspring at PND90 and from 352 to 288 μg/g in Cd-exposed offspring at PND120. GD18 Females: control, n=9; CdCl2, n=11. Males: control, n=8; CdCl2, n=10. PND21 Females: control, n=4; CdCl2, n=4. PND42 Females: control, n=5; CdCl2, n=4. Males: control, n=4; CdCl2, n=4. PND90 Females: control, n=6; CdCl2, n=4. Males: control, n=4; CdCl2, n=6. PND120 Females: control, n=5; CdCl2, n=4. Males: control, n=4; CdCl2, n=5. The level of statistical significance for differences between mean values of control and CdCl2-exposed groups was determined by a two-way ANOVA (treatment, sex) with a Tukey’s post hoc test for all experiments and is indicated by * (p<0.05). All images show mean ± SD.

### 3.4. Gestational Cd mediated changes in metallothionein and metal transporter gene expression

Quantitative RT-PCR was performed on hepatic and placental mRNA to evaluate the progression of dysregulation of mRNA expression of genes involved in metal homeostasis (Table 3 and 4). At GD18, expression of *Mt1* and *Mt2* were increased by Cd exposure in both placenta and liver. In exposed placenta, *Mt1* was 4.8 ± 1.07-fold upregulated in females (p < .0001) and 2.0 ± 0.64 -fold upregulated in males (p = .04). Expression of *Mt2* was significantly increased (p < .0001) only in female placenta (Table 3). Placental mRNA expression of *Slc39a14* was upregulated in females (p < .0001) but not males (p = .19; Table 3). Transcripts encoding the two transporters primarily responsible for efflux of divalent metal cations from the placenta into the cord blood (*Dmt1* and *ZnT2;* Figure 4) were both increased in placenta of Cd exposed females and males (*Dmt1:* females: p < .0001; males: p = .16. *ZnT2:* females: p = .004; males: p = .39; Table 3).

**Table 3.** qRT-PCR Results for mRNA Expression Changes in Placenta and Developing Liver.

| GD18 Placenta |  |  | Female |  |  |  | Male |  |  |  |
| --- | --- | --- | --- | --- | --- | --- | --- | --- | --- | --- |
| Gene | Test statistic | P-value | Fold Change (Mean ± SD) | Sample size (control, CdCl<sub>2</sub>) | Direction | P-value | Fold Change (Mean ± SD) | Sample size (control, CdCl<sub>2</sub>) | Direction | P-value |
| **Slc11a2** | **F(1, 18) = 20.7** | **.0002** | **3.76 ± 1.04** | **6, 6** | **↑** | **<.0001** | 1.86 ± 1.22 | 4, 6 | - | .16 |
| **Slc30a2** | **F(1, 18) = 2.5** | **.01** | **2.2 ± 0.71** | **6, 6** | **↑** | **.004** | 1.35 ± 0.68 | 4, 6 | - | .39 |
| Slc30a10 | F(1, 17) = 1.8 | .19 | 0.59 ± 0.56 | 6, 5 | - | - | 0.78 ± 0.55 | 4, 6 | - | - |
| Slc39a4 | F(1, 18) = 4.6 | .046 | 2.33 ± 1.06 | 6, 6 | - | .007 | 1.08 ± 0.66 | 4, 6 | - | .87 |
| Slc39a8 | F(1, 18) = 3.6 | .07 | 1.54 ± 0.41 | 6, 6 | - | - | 1.15 ± 0.26 | 4, 6 | - | - |
| **Slc39a14** | **F(1, 18) = 19.4** | **.0003** | **2.72 ± 0.76** | **6, 6** | **↑** | **<.0001** | 1.52 ± 0.59 | 4, 6 | - | .19 |
| **Mt1** | **F(1, 17) = 48.7** | **<.0001** | **4.78 ± 1.07** | **6, 6** | **↑** | **<.0001** | **1.95 ± 0.64** | **4, 6** | **↑** | **0.04** |
| **Mt2** | **F(1, 18) = 28.9** | **<.0001** | **2.72 ± 0.53** | **6, 6** | **↑** | **<.0001** | 1.34 ± 0.59 | 4, 6 | - | .25 |
| Mt3 | ND | ND | ND | 6, 6 | - | ND | ND | 4, 6 | - | ND |
| GD18 Liver |  |  | Female |  |  |  | Male |  |  |  |
| Gene | Test statistic | P-value | Fold Change (Mean ± SD) | Sample size (control, CdCl<sub>2</sub>) | Direction | P-value | Fold Change (Mean ± SD) | Sample size (control, CdCl<sub>2</sub>) | Direction | P-value |
| **Mt1** | **F(1, 19) = 36.2** | **<.0001** | **0.34 ± 0.1** | **6, 7** | **↓** | **<.0001** | 0.82 ± 0.23 | 4, 6 | - | .11 |
| Mt2 | F(1, 19) = 1.2 | .29 | 0.81 ± 0.28 | 6, 7 | - | - | 0.93 ± 0.26 | 4, 6 | - | - |
| Mt3 | ND | ND | ND | 6, 7 | - | ND | ND | 4, 6 | - | ND |
| PND1 Female Liver |  |  |  |  |  | PND21 Female Liver |  |  |  |  |
| Gene | Test statistic | Fold Change (Mean ± SD) | Sample size (control, CdCl<sub>2</sub>) | Direction | P-value | Test statistic | Fold Change (Mean ± SD) | Sample size (control, CdCl<sub>2</sub>) | Direction | P-value |
| Slc11a2 | t(1, 6) = 0.84 | 0.90 ± 0.17 | 4, 4 | - | .43 | t(1, 4) = 1.41 | 1.40 ± 0.47 | 3, 3 | - | .23 |
| **Slc25a16** | **t(1, 6) = 3.0** | **1.43 ± 0.24** | **4, 4** | **↑** | **.02** | t(1, 4) = 5.6 | 1.78 ± 0.24 | 3, 3 | - | .005 |
| Slc30a10 | t(1, 6) = 1.7 | 0.75 ± 0.22 | 4, 4 | - | .14 | **t(1, 4) = 3.1** | **2.04 ± 0.57** | **3, 3** | **↑** | **.04** |
| **Slc39a4** | **t(1, 6) = 2.8** | **1.8 ± 0.47** | **4, 4** | **↑** | **.04** | t(1, 4) = 3.0 | 1.87 ± 0.28 | 3, 3 | - | .04 |
| **Slc39a6** | **t(1, 6) = 4.5** | **0.81 ± 0.08** | **4, 4** | **↓** | **.004** | t(1, 4) = 1.4 | 0.72 ± 0.22 | 3, 3 | - | .24 |
| Slc39a8 | t(1, 6) = 0.39 | 0.97 ± 0.04 | 4, 4 | - | .39 | t(1, 4) = 0.12 | 0.97 ± 0.31 | 3, 3 | - | .91 |
| Slc39a14 | t(1, 6) = 0.78 | 1.09 ± 0.10 | 4, 4 | - | .47 | **t(1, 4) = 4.3** | **0.68 ± 0.05** | **3, 3** | **↓** | **.01** |

Values represent mean ± SD fold-change ( $2^{-\Delta\Delta C_t}$ ) for each gene normalized to control. Significant differences are indicated in bold, $p < .05$ . ND; not detected

**Table 4.**
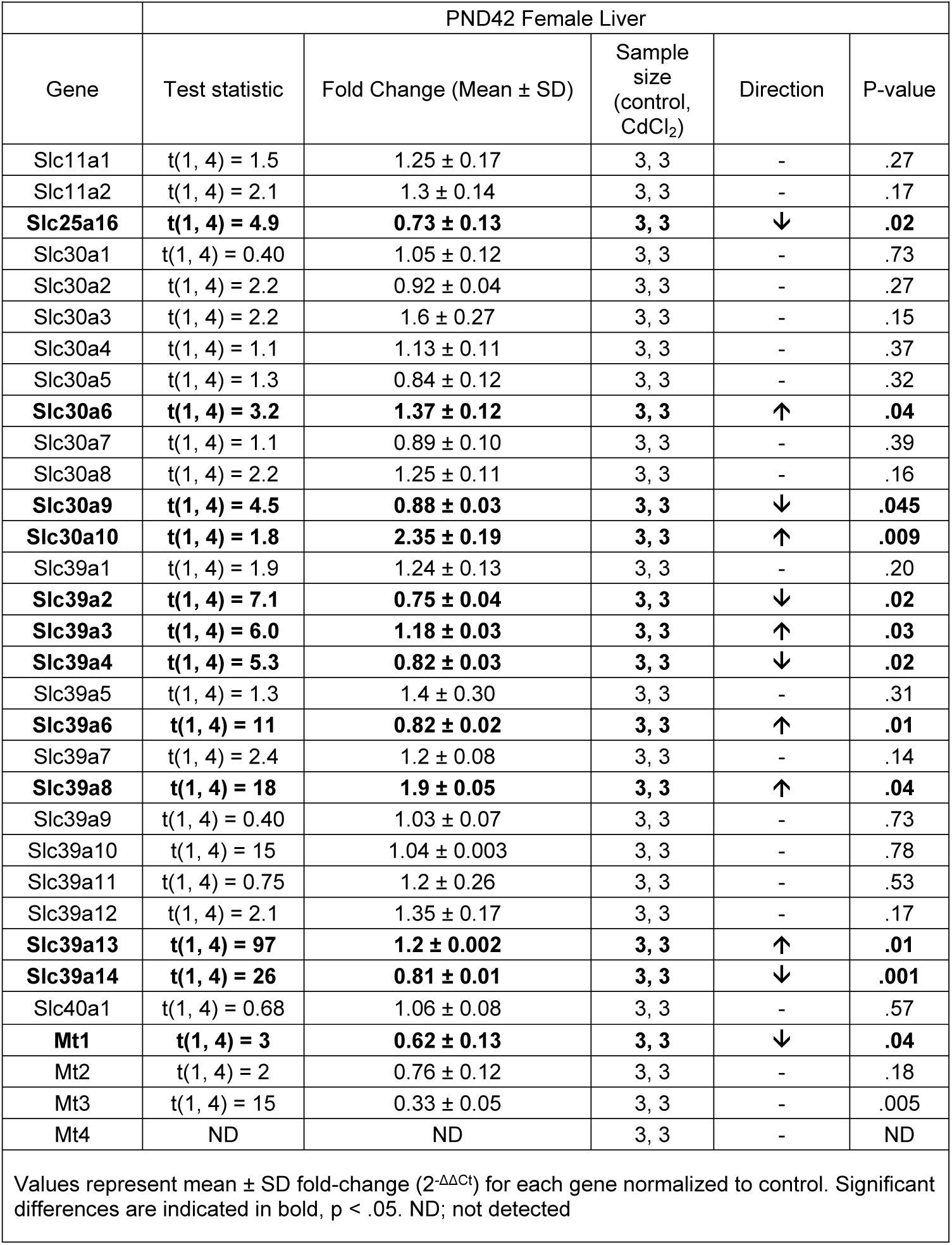
qRT-PCR Results of mRNA Expression Changes in Adult Female Liver.

|  |  |  |  |  |  |
| --- | --- | --- | --- | --- | --- |
| PND42 Female Liver |  |  |  |  |  |
| Gene | Test statistic | Fold Change (Mean ± SD) | Sample size (control, CdCl<sub>2</sub>) | Direction | P-value |
| Slc11a1 | t(1, 4) = 1.5 | 1.25 ± 0.17 | 3, 3 | - | .27 |
| Slc11a2 | t(1, 4) = 2.1 | 1.3 ± 0.14 | 3, 3 | - | .17 |
| **Slc25a16** | **t(1, 4) = 4.9** | **0.73 ± 0.13** | **3, 3** | **↓** | **.02** |
| Slc30a1 | t(1, 4) = 0.40 | 1.05 ± 0.12 | 3, 3 | - | .73 |
| Slc30a2 | t(1, 4) = 2.2 | 0.92 ± 0.04 | 3, 3 | - | .27 |
| Slc30a3 | t(1, 4) = 2.2 | 1.6 ± 0.27 | 3, 3 | - | .15 |
| Slc30a4 | t(1, 4) = 1.1 | 1.13 ± 0.11 | 3, 3 | - | .37 |
| Slc30a5 | t(1, 4) = 1.3 | 0.84 ± 0.12 | 3, 3 | - | .32 |
| **Slc30a6** | **t(1, 4) = 3.2** | **1.37 ± 0.12** | **3, 3** | **↑** | **.04** |
| Slc30a7 | t(1, 4) = 1.1 | 0.89 ± 0.10 | 3, 3 | - | .39 |
| Slc30a8 | t(1, 4) = 2.2 | 1.25 ± 0.11 | 3, 3 | - | .16 |
| **Slc30a9** | **t(1, 4) = 4.5** | **0.88 ± 0.03** | **3, 3** | **↓** | **.045** |
| **Slc30a10** | **t(1, 4) = 1.8** | **2.35 ± 0.19** | **3, 3** | **↑** | **.009** |
| Slc39a1 | t(1, 4) = 1.9 | 1.24 ± 0.13 | 3, 3 | - | .20 |
| **Slc39a2** | **t(1, 4) = 7.1** | **0.75 ± 0.04** | **3, 3** | **↓** | **.02** |
| **Slc39a3** | **t(1, 4) = 6.0** | **1.18 ± 0.03** | **3, 3** | **↑** | **.03** |
| **Slc39a4** | **t(1, 4) = 5.3** | **0.82 ± 0.03** | **3, 3** | **↓** | **.02** |
| Slc39a5 | t(1, 4) = 1.3 | 1.4 ± 0.30 | 3, 3 | - | .31 |
| **Slc39a6** | **t(1, 4) = 11** | **0.82 ± 0.02** | **3, 3** | **↑** | **.01** |
| Slc39a7 | t(1, 4) = 2.4 | 1.2 ± 0.08 | 3, 3 | - | .14 |
| **Slc39a8** | **t(1, 4) = 18** | **1.9 ± 0.05** | **3, 3** | **↑** | **.04** |
| Slc39a9 | t(1, 4) = 0.40 | 1.03 ± 0.07 | 3, 3 | - | .73 |
| Slc39a10 | t(1, 4) = 15 | 1.04 ± 0.003 | 3, 3 | - | .78 |
| Slc39a11 | t(1, 4) = 0.75 | 1.2 ± 0.26 | 3, 3 | - | .53 |
| Slc39a12 | t(1, 4) = 2.1 | 1.35 ± 0.17 | 3, 3 | - | .17 |
| **Slc39a13** | **t(1, 4) = 97** | **1.2 ± 0.002** | **3, 3** | **↑** | **.01** |
| **Slc39a14** | **t(1, 4) = 26** | **0.81 ± 0.01** | **3, 3** | **↓** | **.001** |
| Slc40a1 | t(1, 4) = 0.68 | 1.06 ± 0.08 | 3, 3 | - | .57 |
| **Mt1** | **t(1, 4) = 3** | **0.62 ± 0.13** | **3, 3** | **↓** | **.04** |
| Mt2 | t(1, 4) = 2 | 0.76 ± 0.12 | 3, 3 | - | .18 |
| Mt3 | t(1, 4) = 15 | 0.33 ± 0.05 | 3, 3 | - | .005 |
| Mt4 | ND | ND | 3, 3 | - | ND |
| Values represent mean ± SD fold-change (2<sup>-ΔΔCt</sup>) for each gene normalized to control. Significant differences are indicated in bold, p < .05. ND; not detected |  |  |  |  |  |

**Figure 4.**
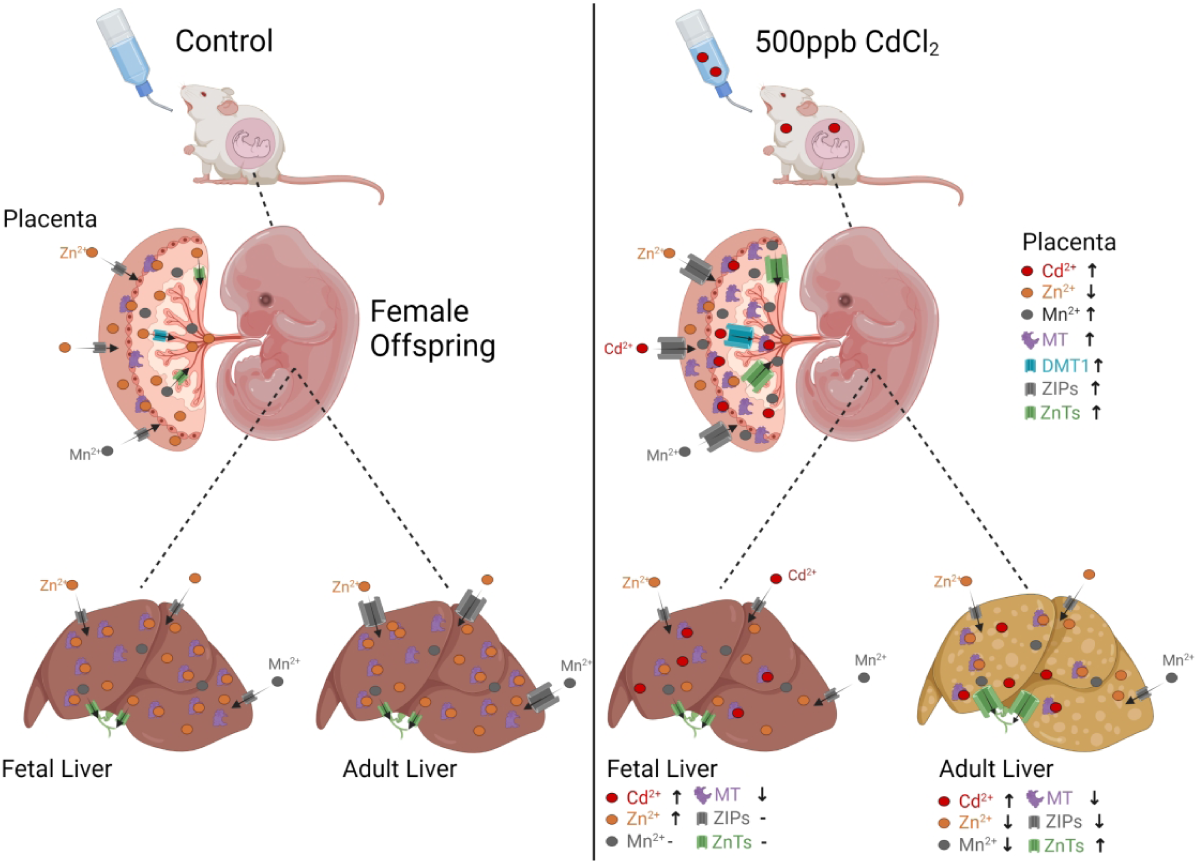
Graphic Representation of Gestational Cd Disruption Placental and Hepatic Metal Concentration and Alters mRNA Expression of Metal Ion Transporters. This graphic illustrates that dams were exposed to 500ppb CdCl_2_ in drinking water. Exposed dams accumulated 850X more Cd than control dams by GD18. At GD18, 15X more Cd was detected in the placenta and ∼8X more Cd in the liver of exposed female offspring than control female offspring. Gestational Cd exposure resulted in decreased placental Zn and increased placental Mn in exposed female offspring. Metallothionein mRNA expression was upregulated in exposed female placenta. The transporter responsible for uptake from maternal blood vessels into the placenta (ZIP14) and the transporters responsible for uptake from the placenta into the fetal cord blood (DMT1, ZnT2) were upregulated in the placentas of exposed female offspring relative to same-sex controls. In fetal livers from Cd-exposed animals, Cd and Zn concentration were increased, whereas metallothionein levels were downregulated and remained downregulated into adulthood. The increased hepatic Zn was transient. Metal ion transporters responsible for influx of divalent metal cations into the liver were significantly downregulated in adulthood, whereas efflux transporters responsible for export into the bile were significantly upregulated. By adulthood, the hepatic Zn phenotypes reversed, with adult hepatic Zn decreases in exposed female offspring.

### 3.5 Postnatal changes in metallothionein and metal transporter gene expression

Because hepatic and metabolic pathology was observed only in Cd-exposed females, analysis of postnatal changes in mRNA expression of genes encoding metal ion transporters and metallothionein was limited to female offspring. At PND1, gestational Cd exposure increased Slc39a4 (p = .04) and decreases Slc39a6 (p = .004) mRNA expression (Table 3). Expression of mRNA zinc transporter associated with mitochondrial defects, *Slc25a16*, was significantly decreased at PND1, decreased expression was observed at all time points (Table 3 and Table 4). Expression of mRNA encoding the primary Zn influx (Slc39a14, *Zip14)* or efflux (Slc30a10, ZnT10) transporters were not significantly changed at PND1. At PND21 and PND42 mRNA expression of *Slc39a14* was significantly decreased and *Slc30a10* expression was significantly (Table 3 and Table 4). At PND42, significant changes in expression of 11 of the 28 metal ion transporter genes analyzed were observed (Table 4). Expression of 4 of 17 influx transporters (e.g., Slc39 family) were significantly decreased, while expression of mRNA encoding 3 transporters was increased (Table 4). At PND42 mRNA expression of primary efflux transporters *Slc30a6* and *Slc30a10* were significantly increased, and Slc30a9 was decreased by gestational Cd exposure. Significant decreases in expression of metallothionein (*Mt1, Mt3)* were also observed at PND42 (Table 4).

### 3.6 Promotor DNA methylation analysis

In females gestationally exposed to Cd, there were no differences in the methylation levels at CpG sites within putative promoter regions upstream of the gene encoding Slc30a10 (Control: 1.6 ± 0.88 %; CdCl_2_: 1.3 ± 0.93 %; p = .42), Slc25a16 (Control: 0.60 ± 0.35 %; CdCl_2_: 0.48 ± 0.40 %; p = .53), Slc39a6 (Control: 0.78 ± 0.42 %; CdCl_2_: 0.45 ± 0.29 %; p = .09), or Slc39a14 (Control: 1.13 ± 1.26 %; CdCl_2_: 2.06 ± 1.58 %; p = .21; Supplemental Table 4). All four of the putative promoter regions upstream of genes encoding metal ion transporters are fully unmethylated. Promoter regions upstream of the gene encoding Mt1, the CpG sites were all approximately 25% methylated with no differences detected between control and Cd-exposed groups (Control: 24 ± 17 %; CdCl_2_: 26 ± 15 %; p = .79; Supplemental Table 4).

## 4. Discussion

The primary study findings were that sex-specific adverse metabolic effects of gestational Cd exposure observed previously were associated with increased transplacental Cd transfer from dam to female offspring, with subsequent Cd accumulation in female liver. It is considered probable that Cd accumulation is programming female offspring for hepatic dysfunction, resulting in the adverse metabolic disruption observed in adult females (Jackson et al. 2020). In our previous study we demonstrated that maternal blood Cd levels of exposed dams at mating was similar to geometric mean blood Cd concentrations for women of childbearing age in the US. Additional evidence for human relevance of this mouse model was demonstrated here with the finding of mean maternal liver concentrations of the Cd-exposed group (364 ± 99 ug/kg) was comparable to nonoccupationally human population in the US (ATSDR 2012 Sep). Overall, our findings that female-specific gestational uptake of Cd by the fetus results in severe metabolic pathology later-in-life and indicates that even small, short-term exposures to Cd during development are harmful and likely a considered significant threat to human metabolic health. As is recognized for lead exposure, there may be no safe level of Cd exposure during development.

### 4.1 Cd accumulates in male and female placenta and decreases efficiency of male placenta

In our previous study, compared to unexposed controls, Cd exposure resulted in a small but significant 5.5 % increase in the female GD18 fetus weight, whereas male fetus weight was unchanged (Jackson et al. 2020). Here, exposed male, but not female GD18 placenta weight was increased by 22% leading to a 15% decline in placental efficiency. Placental efficiency is a proxy measure for placental function and metabolic rate and quantified as the ratio of fetal to placental weight, reflecting grams of fetus produced per grams placenta (Hayward et al. 2016). Observed increases in placenta weight suggest male-specific placental compensatory mechanisms are occurring in response to Cd that warrant additional study. In mammals, Cd was previously demonstrated to poorly cross the placenta; however, previous analyses have not accounted for offspring sex and analyzed Cd concentrations in whole fetal homogenates, thereby limiting sensitivity to detect Cd accumulation (Mikolić et al. 2015). By contrast, here we observed significant Cd levels in GD18 female fetal livers that were 5 times greater than background levels found in males. Placental Cd levels were similarly elevated in both males and females, with mean total Cd of 7.1 ng in females and 7.5 ng in males, which is consistent with previous findings that material Cd accumulates in placenta. In adult offspring, significant Cd was detected only in female offspring liver. Those findings suggest Cd preferentially crosses female placenta in CD-1 mice and accumulates in liver. The retention of Cd in adult liver is consistent with the longer elimination half-life of Cd in females (Taguchi and Suzuki 1981; Satarug et al. 2010). It is probable that Cd accumulated in both placenta and livers of female offspring is exerting direct toxic effects contributing to the sex-specific metabolic disease and obesity in adulthood.

### 4.2 Gestational Cd alters essential metal levels

In addition to previously demonstrated Cd-induced decreases in hemoglobin levels, we observed that Cd exposure significantly decreased fetal and adult liver Fe and placental Zn concentrations and increased placental Mn in both sexes. These findings are consistent with evidence from human birth cohort studies showing Cd exposures are associated anemia, iron and zinc deficiencies (Akesson et al. 2002; Horiguchi et al. 2011; Ikeh-Tawari et al. 2013; Vidal et al. 2015; Luo et al. 2017). Along with anemia, maternal Zn deficiency is associated with fetal death, congenital malformations, intrauterine growth retardation, reduced birth weight, and other adverse health outcomes (Chaffee and King 2012). In rodent models, maternal Zn deficiency can also induce insulin resistance in adult offspring, even with adequate adult Zn (Jou et al. 2010). In the context of those results, placental Zn deficiencies in Cd exposed females may play a programing role in the etiology of Cd-induced adult female metabolic disease.

Along with placental metal concentration changes, sex-specific disruption of essential metal concentrations was evident in GD18 livers. Specifically, Fe was decreased 15% in male liver, and Zn increased 26% in female liver, suggesting differences in compensatory responses to Cd. Impacts were persistent with decreased hepatic Zn, Fe, and Mn levels in Cd exposed adult females, and a significant decrease in Fe in adult males. In females, placenta and liver Cd levels positively correlate with liver Zn, and are negatively correlated with placental Zn, providing further evidence that placental Cd influences female essential metal concentrations. The persistent changes in female divalent cation levels suggest gestational Cd is programming lifelong alterations in hepatic essential metal homeostasis.

### 4.3 Dysregulation of hepatic and placental metal ion transporters and metallothionein

Members of SLC39 and SLC30 family transporters, DMT1, and metallothionein are important regulators of essential metal homeostasis and play critical roles in Cd toxicity. Expression of these transporters is tissue-specifically responsive to Fe, Zn, and Cd (Dufner-Beattie et al. 2003; Taylor et al. 2005; Aydemir and Cousins 2018). However, effects of “low-dose” developmental or chronic Cd exposure on later-in-life expression in metabolically relevant tissues is unknown. Zinc and Cd arrive in placenta from maternal blood via divalent cation transporters and are then delivered to cord blood and fetus primarily via ZnT2, Zip14, and Dmt1 (Espart et al. 2018). During pregnancy, Cd is transported to cord blood through placental DMT1, causing downregulation of Zn transporters, reduced Zn transport to the fetus, resulting in fetal Zn deficiency (Espart et al. 2018). We observed upregulation of *Slc39a14*, which encodes the *Zip14* Zn transporter responsible for uptake from maternal blood vessels into the placenta, ZnT2, and Dmt1 in Cd exposed female placentas, which likely explains female-specific increased hepatic Cd and Zn at GD18 (Figure 4).

In Cd-exposed females, expression of 12 of 28 analyzed metal ion transporter genes were modified in adult liver, including primary influx transporter, *Slc39a14*, and primary efflux transporter, *Slc30a10*. Additionally, gestational Cd exposure downregulated expression of metal transporters at PND1, PND21, and PND42 including *Slc39a4 (Zip4)*, and *Slc25a16* (Jackson et al. 2020). *Zip4* encodes a tissue-specific, zinc-regulated zinc transporter in mice inversely responsive to dietary Zn levels (Dufner-Beattie et al. 2003). Heterozygous *Zip4*-knockout mice display varied phenotypes including severe growth retardation exacerbated by maternal Zn deficiency and ameliorated by maternal Zn supplementation (Dufner-Beattie et al. 2007). Slc25a16 is associated with mitochondrial defects and reduction in mitochondrial coenzyme A levels (Prohl et al. 2001; Gutiérrez-Aguilar and Baines 2013). Based on observed effects, gestational Cd appears to act as a Zn mimic, inducing a long-term epigenetic alteration in metal transporter expression that persists into adulthood and contributes to fatty liver disease and insulin insensitivity pathogenesis.

Gestational Cd exposure also altered expression of placental and liver metallothionein mRNA. The metal-binding metallothioneins are involved in regulation of absorption, transport, and homeostasis of essential trace metals, and have cytoprotective functions binding and sequestering toxic metals including Cd (Palmiter 1987; Andrews 1990; Dalton et al. 1994; Klaassen et al. 2009; Sabolić et al. 2010). *In vivo* and *in vitro* experimental studies demonstrated Cd-induced changes in metal ion transporter and metallothionein expression are involved in the mechanisms of Cd-induced toxicity and dysregulation of Zn homeostasis (Akesson et al. 2002; Sabolić et al. 2010; Thévenod 2010; Horiguchi et al. 2011; Genchi et al. 2020). Cumulative Cd toxicity results from high-affinity binding of Cd^2+^ by metallothionein and a resulting dose-dependent loss of protective antioxidant mechanisms leading to oxidative damage (Sabolić et al. 2010; Nair et al. 2013; Nemmiche 2016 Nov 1). Metallothionein expression is induced by increased free Zn^2+^ levels via metal-response element binding transcription factor 1 (Radtke et al. 1993: 1; Andrews 2001: 1; Bhandari et al. 2017) and by free Cd^2+^ indirectly through release of metallothionein-bound Zn^2+^ (Kägi and Schäffer 1988; Klaassen et al. 2009). Cadmium-induced alterations in metallothionein expression contribute to disruptions of essential metal homeostasis. Both major metallothionein isoforms, *Mt1* and *Mt2*, were upregulated in placenta of Cd exposed females, consistent with previous work demonstrating placental induction of metallothionein in rats gestationally exposed to Cd (Nakamura et al. 2012). In males, only *Mt1* mRNA expression was significantly upregulated. Paradoxically, hepatic metallothionein was downregulated at GD18 and PND42 in females, further indicating disrupted metal sensing is a more complex interplay during gestation than free metal ions directly induing metallothionein production. This is supported by studies in transgenic mice overexpressing MT1 that accumulate Zn in maternal compartments and are resistant to teratogenic effects of Zn deficiency during pregnancy, whereas *Mt1* knockout mice accumulate less Zn in maternal tissues and are susceptible to teratogenic effects of dietary Zn deficiency (Dalton et al. 1996; Dalton et al. 1997; Bittel et al. 1998; Andrews and Geiser 1999). The observed metallothionein downregulation may contribute to decreased Zn accumulation in female liver and adverse metabolic impacts resulting from gestational Cd exposure in females.

### 4.4 Gestational Cd did not alter CpG methylation of transporter or metallothionein promoters

Due to early and persistent disruption of mRNA expression of metal ion transporters and metallothionein, gestational Cd seemingly causes lifelong epigenetic expression changes. However, differences in CpG DNA methylation within putative promoter regions upstream of Zip14, ZnT10, Zip6, Slc25a16, or the primary hepatic metallothionein Mt1 were not observed. The CpG island in metal ion transporter promoters were essentially unmethylated and unchanged by Cd exposure. Epigenetic modifications are one type of mechanism representing an organismal response to environmental stressors, yet regions of the epigenome responsive to Cd or other trace metals are largely unknown (Heijmans et al. 2009). Because both MT isoforms are similarly responsive to induction by Cd exposure, the two genes likely share common promoter elements (Andersen et al. 1983; Searle et al. 1984). Demethylation of DNA sequences in the vicinity of the MT1 gene correlates with upregulation of mRNA expression (Compere and Palmiter 1981). In the promoter region upstream of the mouse *Mt1* gene, no differences were detected in CPG methylation at any CpG site examined. Because CpG methylation upstream of genes encoding metal ion transporters and metallothionein were unchanged, transcriptional epigenetic regulation appears not to involve promoter proximal regulation of CpG methylation. Considering the absence of changes in DNA methylation, the notable Cd deposition specifically in female livers may itself be the mechanistic drive of gestational Cd-induced adult disease. Early accumulation of Cd in female livers may directly program adult disease via developmental toxicity that alters lifelong metabolic function. This interpretation is consistent with lifelong changes in metallothionein and metal ion transporter mRNA expression leading to decreased hepatic Zn, Fe, and Mn in adult females.

## 5. Conclusion

Maternal exposure to 500 ppb CdCl_2_ in mice disrupted essential metal concentrations in female offspring. To the best of our knowledge, the finding of detectable Cd levels in female fetal and adult livers following gestational Cd exposure is the first report of an important sex difference in maternal-fetal transfer of Cd. The observed sex-specific fetal transfer and long retention of Cd in female livers was linked to adverse decreases in hepatic Zn concentrations and altered metal ion transporter and metallothionein mRNA expression. These findings suggest gestational Cd accumulation is programming adult metabolic disease through altered essential metal homeostasis.

---

**Supplemental Table 1.**
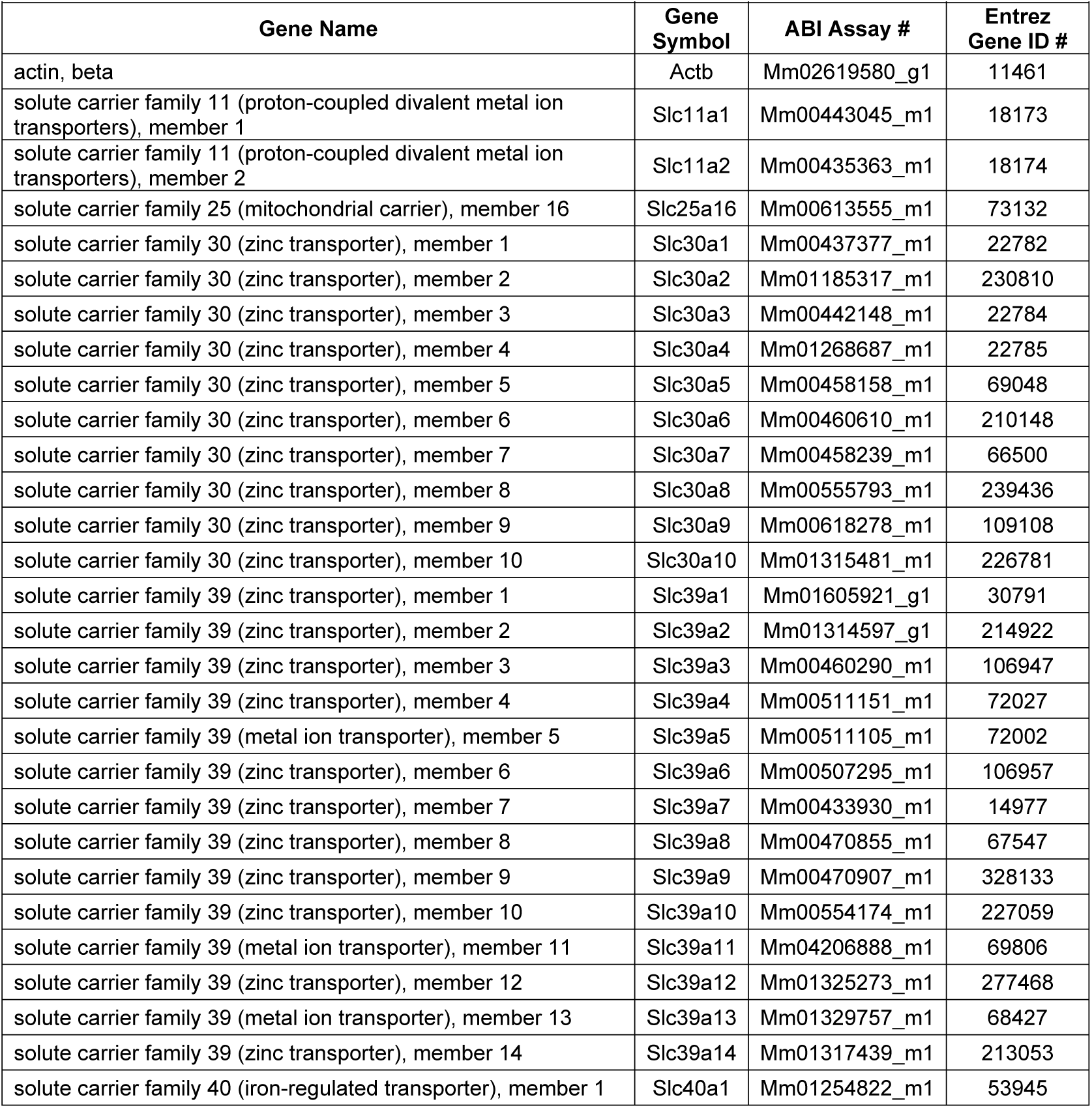
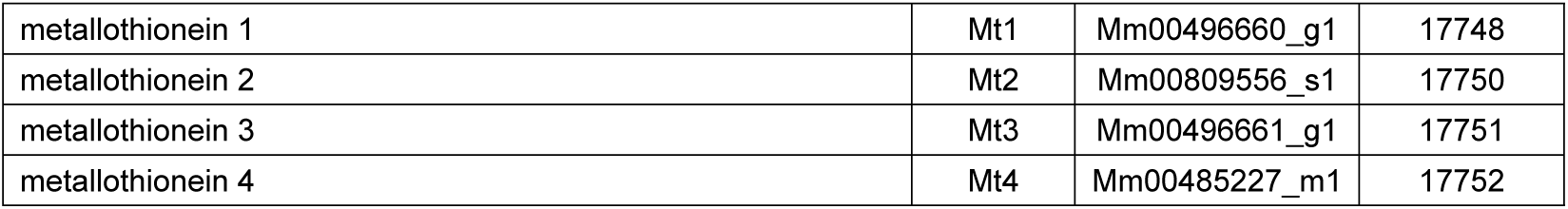
Gene list for qRT-PCR analyses.

| Supplemental Table 1. Gene list for qRT-PCR analyses. |  |  |  |
| --- | --- | --- | --- |
| Gene Name | Gene Symbol | ABI Assay # | Entrez Gene ID # |
| actin, beta | Actb | Mm02619580_g1 | 11461 |
| solute carrier family 11 (proton-coupled divalent metal ion transporters), member 1 | Slc11a1 | Mm00443045_m1 | 18173 |
| solute carrier family 11 (proton-coupled divalent metal ion transporters), member 2 | Slc11a2 | Mm00435363_m1 | 18174 |
| solute carrier family 25 (mitochondrial carrier), member 16 | Slc25a16 | Mm00613555_m1 | 73132 |
| solute carrier family 30 (zinc transporter), member 1 | Slc30a1 | Mm00437377_m1 | 22782 |
| solute carrier family 30 (zinc transporter), member 2 | Slc30a2 | Mm01185317_m1 | 230810 |
| solute carrier family 30 (zinc transporter), member 3 | Slc30a3 | Mm00442148_m1 | 22784 |
| solute carrier family 30 (zinc transporter), member 4 | Slc30a4 | Mm01268687_m1 | 22785 |
| solute carrier family 30 (zinc transporter), member 5 | Slc30a5 | Mm00458158_m1 | 69048 |
| solute carrier family 30 (zinc transporter), member 6 | Slc30a6 | Mm00460610_m1 | 210148 |
| solute carrier family 30 (zinc transporter), member 7 | Slc30a7 | Mm00458239_m1 | 66500 |
| solute carrier family 30 (zinc transporter), member 8 | Slc30a8 | Mm00555793_m1 | 239436 |
| solute carrier family 30 (zinc transporter), member 9 | Slc30a9 | Mm00618278_m1 | 109108 |
| solute carrier family 30 (zinc transporter), member 10 | Slc30a10 | Mm01315481_m1 | 226781 |
| solute carrier family 39 (zinc transporter), member 1 | Slc39a1 | Mm01605921_g1 | 30791 |
| solute carrier family 39 (zinc transporter), member 2 | Slc39a2 | Mm01314597_g1 | 214922 |
| solute carrier family 39 (zinc transporter), member 3 | Slc39a3 | Mm00460290_m1 | 106947 |
| solute carrier family 39 (zinc transporter), member 4 | Slc39a4 | Mm00511151_m1 | 72027 |
| solute carrier family 39 (metal ion transporter), member 5 | Slc39a5 | Mm00511105_m1 | 72002 |
| solute carrier family 39 (zinc transporter), member 6 | Slc39a6 | Mm00507295_m1 | 106957 |
| solute carrier family 39 (zinc transporter), member 7 | Slc39a7 | Mm00433930_m1 | 14977 |
| solute carrier family 39 (zinc transporter), member 8 | Slc39a8 | Mm00470855_m1 | 67547 |
| solute carrier family 39 (zinc transporter), member 9 | Slc39a9 | Mm00470907_m1 | 328133 |
| solute carrier family 39 (zinc transporter), member 10 | Slc39a10 | Mm00554174_m1 | 227059 |
| solute carrier family 39 (metal ion transporter), member 11 | Slc39a11 | Mm04206888_m1 | 69806 |
| solute carrier family 39 (zinc transporter), member 12 | Slc39a12 | Mm01325273_m1 | 277468 |
| solute carrier family 39 (metal ion transporter), member 13 | Slc39a13 | Mm01329757_m1 | 68427 |
| solute carrier family 39 (zinc transporter), member 14 | Slc39a14 | Mm01317439_m1 | 213053 |
| solute carrier family 40 (iron-regulated transporter), member 1 | Slc40a1 | Mm01254822_m1 | 53945 |
| metallothionein 1 | Mt1 | Mm00496660_g1 | 17748 |
| metallothionein 2 | Mt2 | Mm00809556_s1 | 17750 |
| metallothionein 3 | Mt3 | Mm00496661_g1 | 17751 |
| metallothionein 4 | Mt4 | Mm00485227_m1 | 17752 |

**Supplemental Table 2.**
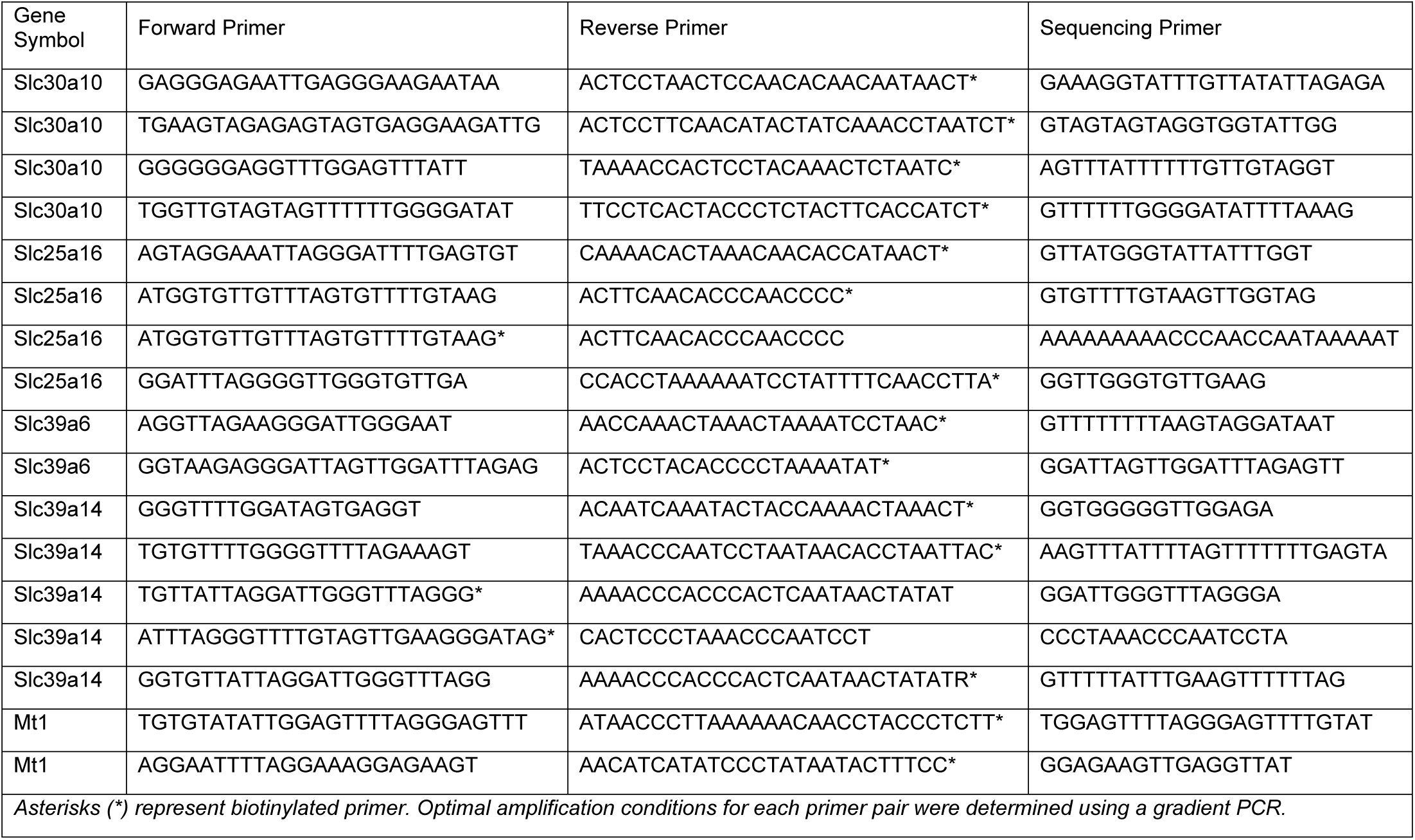
PCR and pyrosequencing primers (5’ - 3’)

**Supplemental Table 3.**
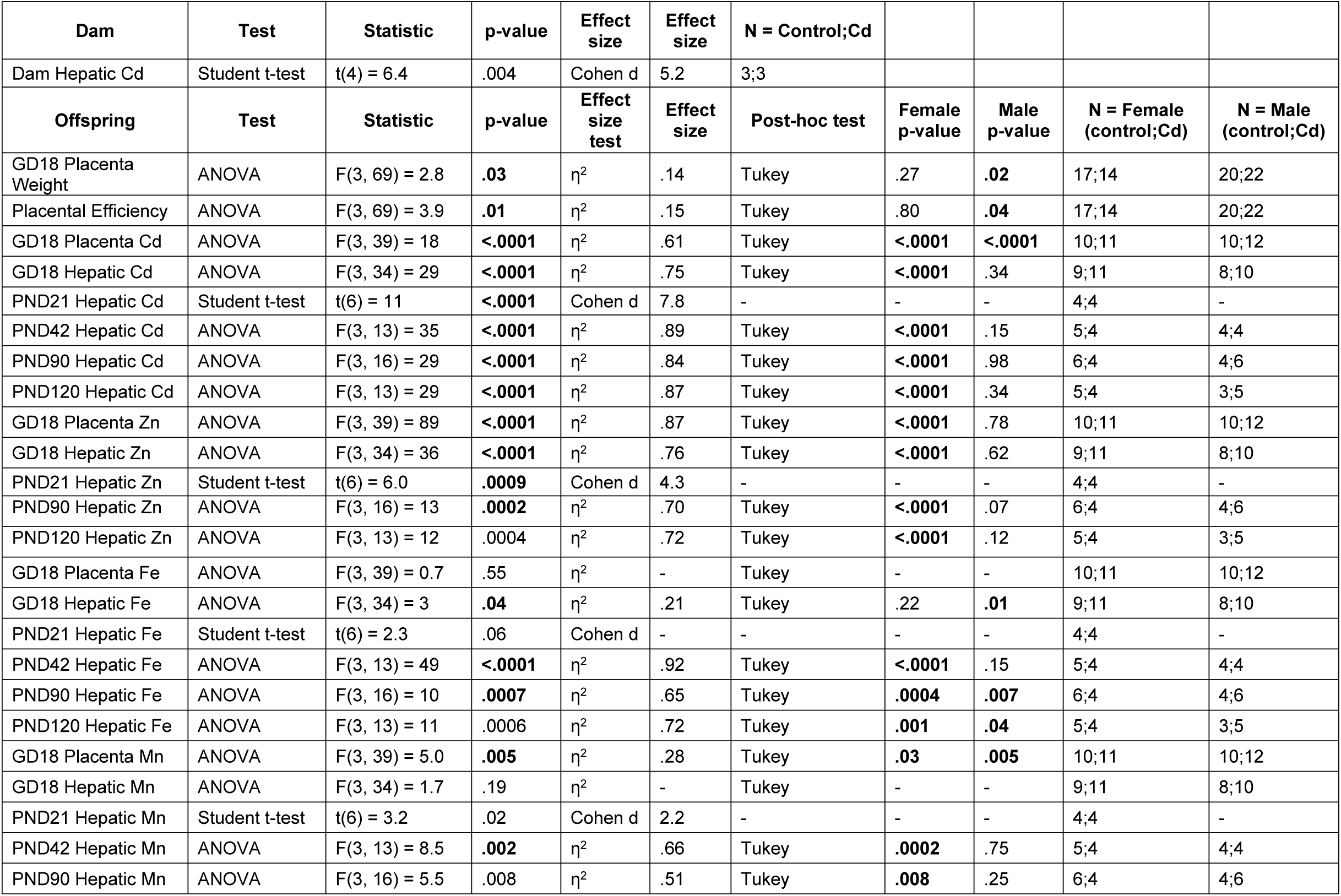
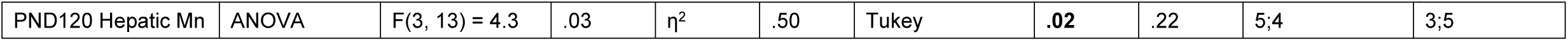
Statistical summary.

| Dam | Test | Statistic | p-value | Effect size | Effect size | N = Control;Cd |  |  |  |  |
| --- | --- | --- | --- | --- | --- | --- | --- | --- | --- | --- |
| Dam Hepatic Cd | Student t-test | t(4) = 6.4 | .004 | Cohen d | 5.2 | 3;3 |  |  |  |  |
| Offspring | Test | Statistic | p-value | Effect size test | Effect size | Post-hoc test | Female p-value | Male p-value | N = Female (control;Cd) | N = Male (control;Cd) |
| GD18 Placenta Weight | ANOVA | F(3, 69) = 2.8 | .03 | $\eta^2$ | .14 | Tukey | .27 | .02 | 17;14 | 20;22 |
| Placental Efficiency | ANOVA | F(3, 69) = 3.9 | .01 | $\eta^2$ | .15 | Tukey | .80 | .04 | 17;14 | 20;22 |
| GD18 Placenta Cd | ANOVA | F(3, 39) = 18 | <.0001 | $\eta^2$ | .61 | Tukey | <.0001 | <.0001 | 10;11 | 10;12 |
| GD18 Hepatic Cd | ANOVA | F(3, 34) = 29 | <.0001 | $\eta^2$ | .75 | Tukey | <.0001 | .34 | 9;11 | 8;10 |
| PND21 Hepatic Cd | Student t-test | t(6) = 11 | <.0001 | Cohen d | 7.8 | - | - | - | 4;4 | - |
| PND42 Hepatic Cd | ANOVA | F(3, 13) = 35 | <.0001 | $\eta^2$ | .89 | Tukey | <.0001 | .15 | 5;4 | 4;4 |
| PND90 Hepatic Cd | ANOVA | F(3, 16) = 29 | <.0001 | $\eta^2$ | .84 | Tukey | <.0001 | .98 | 6;4 | 4;6 |
| PND120 Hepatic Cd | ANOVA | F(3, 13) = 29 | <.0001 | $\eta^2$ | .87 | Tukey | <.0001 | .34 | 5;4 | 3;5 |
| GD18 Placenta Zn | ANOVA | F(3, 39) = 89 | <.0001 | $\eta^2$ | .87 | Tukey | <.0001 | .78 | 10;11 | 10;12 |
| GD18 Hepatic Zn | ANOVA | F(3, 34) = 36 | <.0001 | $\eta^2$ | .76 | Tukey | <.0001 | .62 | 9;11 | 8;10 |
| PND21 Hepatic Zn | Student t-test | t(6) = 6.0 | .0009 | Cohen d | 4.3 | - | - | - | 4;4 | - |
| PND90 Hepatic Zn | ANOVA | F(3, 16) = 13 | .0002 | $\eta^2$ | .70 | Tukey | <.0001 | .07 | 6;4 | 4;6 |
| PND120 Hepatic Zn | ANOVA | F(3, 13) = 12 | .0004 | $\eta^2$ | .72 | Tukey | <.0001 | .12 | 5;4 | 3;5 |
| GD18 Placenta Fe | ANOVA | F(3, 39) = 0.7 | .55 | $\eta^2$ | - | Tukey | - | - | 10;11 | 10;12 |
| GD18 Hepatic Fe | ANOVA | F(3, 34) = 3 | .04 | $\eta^2$ | .21 | Tukey | .22 | .01 | 9;11 | 8;10 |
| PND21 Hepatic Fe | Student t-test | t(6) = 2.3 | .06 | Cohen d | - | - | - | - | 4;4 | - |
| PND42 Hepatic Fe | ANOVA | F(3, 13) = 49 | <.0001 | $\eta^2$ | .92 | Tukey | <.0001 | .15 | 5;4 | 4;4 |
| PND90 Hepatic Fe | ANOVA | F(3, 16) = 10 | .0007 | $\eta^2$ | .65 | Tukey | .0004 | .007 | 6;4 | 4;6 |
| PND120 Hepatic Fe | ANOVA | F(3, 13) = 11 | .0006 | $\eta^2$ | .72 | Tukey | .001 | .04 | 5;4 | 3;5 |
| GD18 Placenta Mn | ANOVA | F(3, 39) = 5.0 | .005 | $\eta^2$ | .28 | Tukey | .03 | .005 | 10;11 | 10;12 |
| GD18 Hepatic Mn | ANOVA | F(3, 34) = 1.7 | .19 | $\eta^2$ | - | Tukey | - | - | 9;11 | 8;10 |
| PND21 Hepatic Mn | Student t-test | t(6) = 3.2 | .02 | Cohen d | 2.2 | - | - | - | 4;4 | - |
| PND42 Hepatic Mn | ANOVA | F(3, 13) = 8.5 | .002 | $\eta^2$ | .66 | Tukey | .0002 | .75 | 5;4 | 4;4 |
| PND90 Hepatic Mn | ANOVA | F(3, 16) = 5.5 | .008 | $\eta^2$ | .51 | Tukey | .008 | .25 | 6;4 | 4;6 |
| PND120 Hepatic Mn | ANOVA | F(3, 13) = 4.3 | .03 | $\eta^2$ | .50 | Tukey | .02 | .22 | 5;4 | 3;5 |

**Supplemental Table 4.** CpG methylation of putative promoter regions upstream of metallothionein and metal ion transporter genes in female CD-1 mice

| Gene | Control | CdCl<sub>2</sub> | # CpG Sites |
| --- | --- | --- | --- |
| Slc30a10 | 1.64 ± 0.88 (8) | 1.26 ± 0.93 (8) | 15 |
| Slc25a16 | 0.60 ± 0.35 (8) | 0.48 ± 0.40 (8) | 27 |
| Slc39a6 | 0.78 ± 0.42 (8) | 0.45 ± 0.29 (8) | 12 |
| Slc39a14 | 1.13 ± 1.26 (8) | 2.06 ± 1.58 (8) | 14 |
| Mt1 | 23.8 ± 17.0 (7) | 26.2 ± 14.8 (6) | 12 |

Values represent group mean (%) ± SEM; the number of samples analyzed is shown in parenthesis. \* $p < .05$ . CdCl<sub>2</sub> = 500ppb gestational Cadmium Chloride

